# A Network Architecture for Bidirectional Neurovascular Coupling in Rat Whisker Barrel Cortex

**DOI:** 10.1101/602680

**Authors:** Bhadra S. Kumar, Aditi Khot, Srinivasa V Chakravarthy, S Pushpavanam

## Abstract

The neurovascular coupling is mostly considered as a master-slave relationship between the neurons and the cerebral vessels: the neurons demand energy which the vessels supply in the form of glucose and oxygen. In the recent past both theoretical and experimental studies have suggested that the neurovascular coupling is a bidirectional system, a loop that includes a feedback signal from the vessels influencing neural firing and plasticity. An integrated model of bidirectionally connected neural network and vascular network is hence required to understand the relationship between the informational and metabolic aspects of neural dynamics. In this study, we present a computational model of the bidirectional neurovascular system in the whisker barrel cortex and study the effect of such coupling on neural activity and plasticity as manifest in the map formation. In this model, a biologically plausible self-organizing network model of rate coded, dynamic neurons is nourished by a network of vessels modeled using the biophysical properties of blood vessels. The neural layer which is designed to simulate the whisker barrel cortex of rat transmits the vasodilatory signals to the vessels. The feedback from the vessels is in the form of available oxygen for oxidative metabolism whose end result is the ATP necessary to fuel neural firing. The model captures the effect of the feedback from the vascular network on the neuronal map formation in the whisker barrel model under normal and pathological (Hypoxia and Hypoxia-Ischemia) conditions.

**Author Summary:** Although neurovascular coupling is typically depicted as a unidirectional influence originating from the neurons and acting on the cerebral vessels, in reality it forms a bidirectional system, consisting of neuronal energy demand signals transmitted to the vessels, and a feedback of metabolic substrates from the vessels, that influence the neural firing and plasticity. We present a computational model of the bidirectional neurovascular coupling in the whisker barrel cortex of rats and study the effect of such coupling on neural activity and plasticity as manifest in the map formation. The model consists of a biologically plausible self-organizing dynamic, neural network model and a biophysical vascular network model that nourishes the neural network in the form of oxygen necessary for neural oxidative metabolism. The model reproduces the spatio-temporal hemodynamic responses observed in rat whisker barrel cortex. It also demonstrates the essentiality of the vascular feedback on the whisker barrel map formation under normal and pathological conditions.

## Introduction

The brain is an energy-intensive organ in the human body, consuming around 20% of the total cardiac output even though it constitutes just 2% of the total body weight. Despite the high energy demands of this organ, there seems to be no significant energy reservoir in the neural tissue and the energy is provided in a ‘supply on demand’ fashion by an increase in blood flow in proximal blood vessels (functional hyperemia) as a response to neural activity. Classical accounts of neurovascular coupling describe the interaction between neurons and cerebral micro-vessels as a unidirectional influence from neurons to vessels. But the dependence of the neurons on the energy substrates delivered by the vessels raises the possibility of a reverse signal emanating from the vessels and influencing neural dynamics.

The possibility of a retrograde signal from the vessels to the neurons was first discussed under the name of the hemoneural hypothesis by Moore and Cao(1). Sirotin and Das(2) showed that under certain conditions the vascular response precedes the neural response. Leithner and Royl(3) demonstrated a decorrelation between blood volume changes and neural oxygen demand. Such studies strengthen the case for a revision of the notion of a master-slave relationship between neurons and cerebral vessels. The study by Filosa and colleagues (4,5) suggested that vessels can have a direct influence on the neurons not necessarily by releasing energy substrates, but by other mechanisms like, for example, the mechanical pressure exerted by the vessels transduced into electrical signals in the neurons.

The above selection of experimental studies motivates an intriguing question: “What are the vascular influences, if any, on neural performance?” The possibility of a retrograde influence of the cerebral vessels on the neurons was also explored from a computational modeling perspective. Gandrakota et al (Gandrakota, Chakravarthy, and Pradhan, 2010) presented a model of the neuro-glio-vascular network as a bidirectional associative memory with information flowing both in neurovascular and vasculoneural directions. A detailed biophysical model of a single neuron-astrocyte-vessel system by Chander and Chakravarthy(7) showed how the slow vasomotor rhythms can modulate neural firing, thereby reinforcing the idea of a retrograde signal. This model was later simplified and extended to a network level by Chabbria et al(8) where the vascular and glial structures were clubbed to a single gliovascular unit whose output determines the energy level of neurons. In a neurovascular network model proposed by Philips et al(9), desynchronized vascular rhythms were found to improve the efficiency of learning in a neural network trained as an autoencoder. Among the aforementioned set of models, there are either detailed biophysical models at the single unit level or abstract models at a network level. There is a need to create detailed neurovascular models that explore, preferably at a network level, the consequences of a retrograde signal from the vessels to the neurons on neuronal activity and plasticity. This last objective forms the motivation of the present study. In this paper, we develop a network level architecture for neurovascular interaction where the activity of neuron not only influences the vascular activity in the forward direction, but also depends on the feedback from the proximal vessels based on their volume and oxygen saturation.

We have selected the rat whisker barrel cortex as the model system for which we develop a neurovascular network model. Devor et al have studied the rat barrel cortex (10–12) where they found that the neural and hemodynamic response have a nonlinear relationship. Dunn et al (13) found that the spatial extents of metabolic variables in neural tissues and hemodynamic variables in nearby vessels are comparable. Although these experiments qualitatively yielded the same results, the quantitative results seemed to vary widely depending on the presence, absence or type of anesthesia given to the animal. The experiments by Martin et al(14), and Gao et al (15) show that neurovascular coupling in the animal is weaker when it is anesthetized, with an increased latency, which is again a serious challenge.

In order to simulate the vascular influence on neural activity and plasticity, a network level model of neural layer and a network of vascular structure that perfuses this neural layer at the level of capillaries are required. The vascular network which perfuses the whisker barrel cortex layer L4 is modeled using a three-dimensional vascular tree architecture which branches and penetrates the neural layer at the level of capillaries. The anatomy of the vascular network is similar to that described by Blinder et al (16) while the vascular dynamics is similar to that of the Vascular anatomical network (VAN) model of Boas et al(17). The pressure-volume relationship in the blood vessel is assumed to be linear with the linearity coefficient modifiable by neural activity. The activity of a neuron influences the vessels in its neighborhood, while the neuron, in turn, is nourished by the vessels in a neighborhood thereby making the neurovascular network bidirectional (Fig.1).

**Figure 1:**
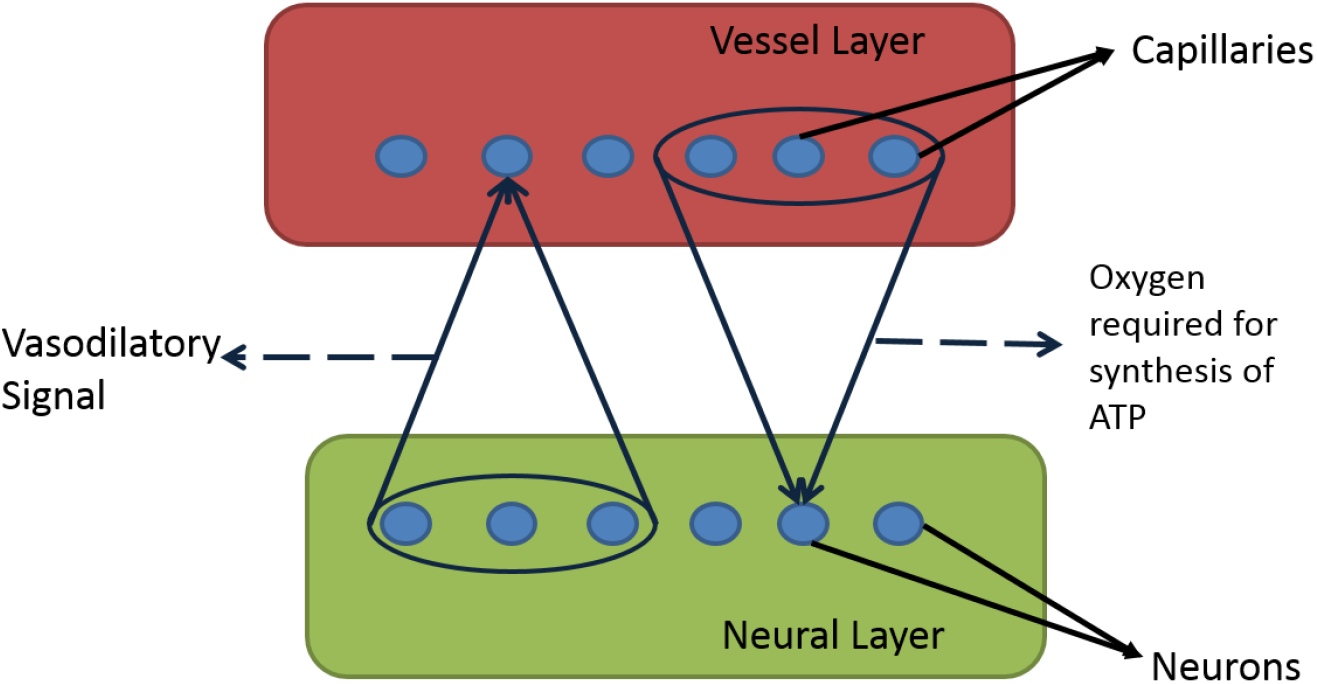
Bidirectional connection between neural and vascular layer. Neurons sends afferent vasodilatory signals to vessels and vessels send oxygen back to neural layer for oxidative phosphorylation

The vasodilation is simplified greatly by using a single variable to represent all possible vasodilatory pathways. The model also allows control of arteriolar dilation and capillary dilation independently which would help to incorporate the studies which show that while the arteriolar dilation is mediated by direct neural influence, the capillary dilation is mostly mediated by astrocytes (18). This kind of network model with modified parameters which simulate hemodynamic response is advantageous as it allows various factors which affect the hemodynamic response characteristics like anesthesia to be easily incorporated and the model could be adapted to simulate different experimental paradigms. The neural dynamics is captured by a laterally interconnecting synergistically self-organizing map (LISSOM) which is a simple yet biologically plausible model to simulate cortical neural activity (19). The dynamics of the LISSOM model basically does a feature extraction similar to the principal component analysis and the response finally forms topographic maps with similar features coded by adjacent neurons. A well-trained LISSOM network evolves to resemble the maps of the rat whisker barrel cortex.

The parameters of the model are optimized to simulate the experiment done by Devor et al(11). The spatial and temporal characteristics of the hemodynamic response in terms of the volume and oxygen saturation agree with the experimental observation. The model is also able to capture the nonlinearity of hemodynamic response. To explore the possibility of studying the vascular influence on neural layer properties, retraining of the neural layer is carried out under various vascular conditions. The results are in agreement with the findings of Ranasinghe et al(20).

## Results

### Neural layer

Fig. 2.b shows the response map of the trained LISSOM network. Each colored blob represents the area in the rat barrel cortex that responds to one particular whisker. The colormap has values from 1 to 24 (representing all 24 whiskers shown in Fig.2.a) which indicates the neurons that respond maximally when the corresponding whisker is stimulated. For example, the neurons coded with number 1 correspond to whisker located at row A and column 0. The studies on whisker barrel cortex show a parabolic arrangement of barrels(21). By imposing an area constraint on the LISSOM sheet, we are able to reproduce the parabolic whisker barrel map seen experimentally.

**Figure 2:**
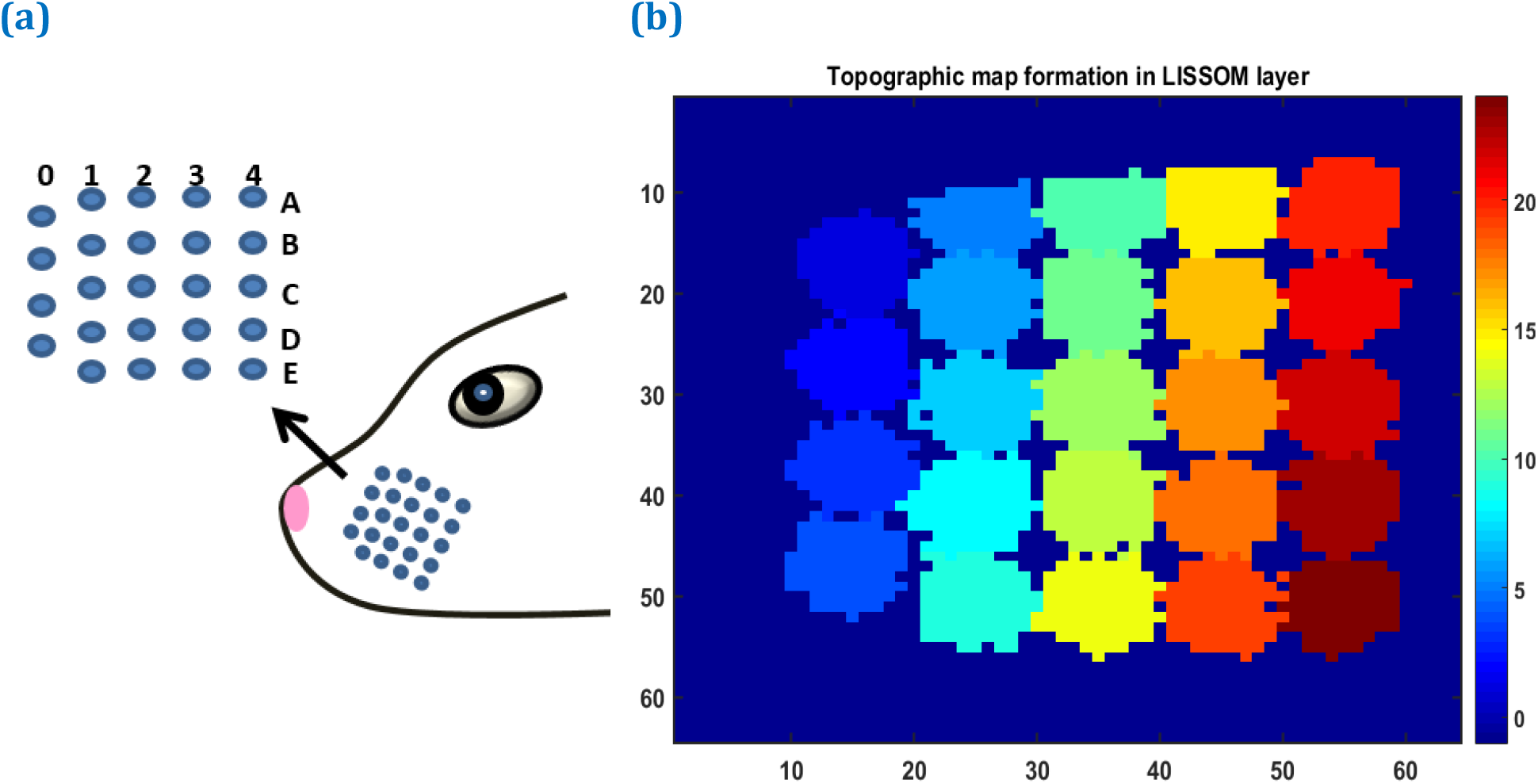
(a) The whisker arrangement on rat whisker pad. (b) The topographic map formed in whisker barrel cortex. The whiskers are given numbers 1-24 such that, column 0 has whiskers 1 to 4; column 1 (5 to 9); column 2 (10 to 14); column 3 (15 to 19); column 4 (20 to 24). In this sheet of 64×64 neurons, each neuron is colour coded to the index of the whisker to which it responds maximally. The nearby neurons respond to the same whisker forming barrels as can be seen in the figure.

### The hemodynamic response following stimulation

Hemodynamic response is observed after giving a brief stimulus of 1s duration to the C3 whisker. It is characterized by the change in total haemoglobin (HbT) which is analogous to the change in volume, change in oxygenated haemoglobin (HbO) and change in deoxygenated haemoglobin (Hb). The variation of HbT with time is shown in Fig. 3.a. The red arrow on the figure indicates the time of presentation of stimulus. As expected, the capillary area corresponding to C3 whisker barrel area shows a response to the input stimulus by increase in HbT which also indicates increase in volume or in other words, vasodilation. The vasodilation begins after a delay and this delay is optimized to match the experimental observation by Devor et al (10). The change in HbT reaches a maximum at around 1.8s post stimulus presentation (22) and decays slowly since the duration of input stimulus is only 1s. Our model also captures the initial dip of HbO (Fig.3.b) at around 0.6s post stimulus presentation as observed by Devor et al and Berwick et al (10,23). HbO reaches its peak at around 1.8s to 2s post stimulus presentation. Hb being complementary to HbO also shows an initial increase around 0.6s after the presentation of the stimulus followed by a steady decrease (Fig.3.c). The HbO then slowly increases when HbT begins to increase and Hb slowly decreases reflecting the hemodynamic response observed in rat whisker barrel cortex by Devor et al (10). Similar results are observed when input stimulus is given to any other whisker. Any whisker stimulation results in a similar hemodynamic response pattern around an area in capillary bed corresponding to the neural area in whisker barrel cortex that responds to the whisker which is stimulated.

**Figure 3:**
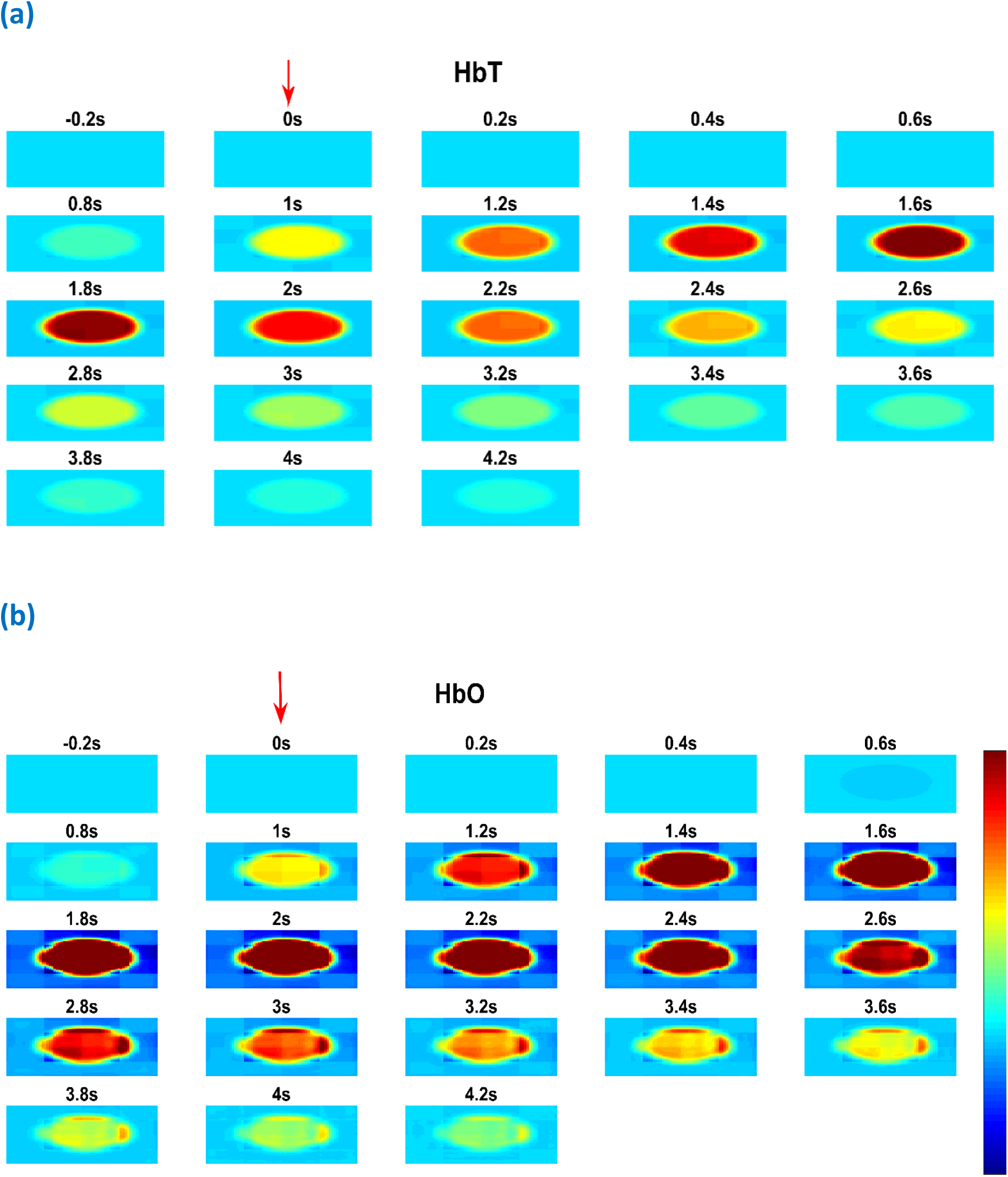
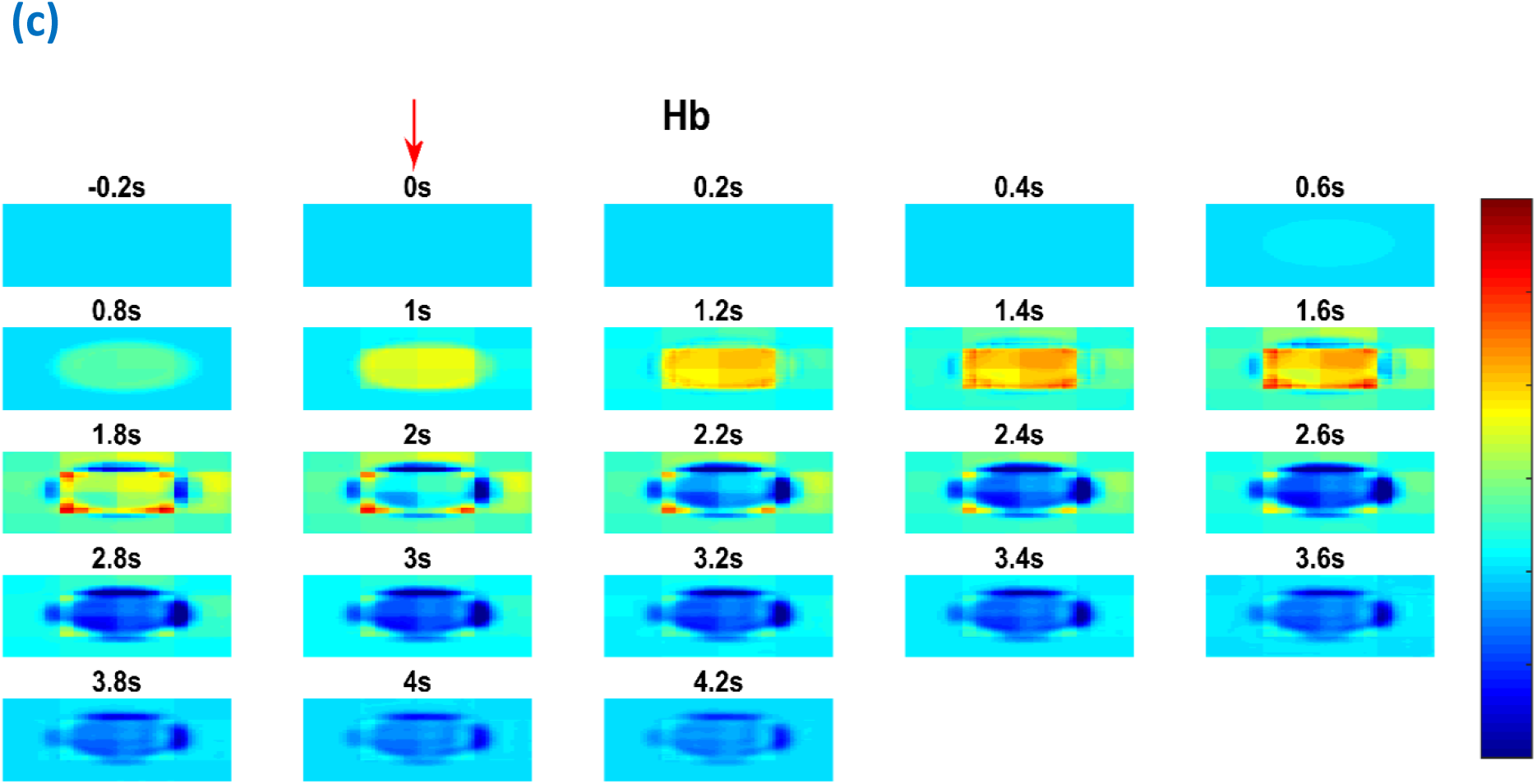
Each plot shows time profile of different hemodynamic variables. The red arrow shows the time of presentation of stimulus. The colorbar on the right shows the percentage change in value (a) The time profile of HbT in the whisker barrel cortex, HbT is maximum at 1.6s post stimulus presentation (b) The time profile of HbO in the whisker barrel cortex. Soon after the stimulus, a dip in HbO can be observed at 0.6s. HbO peaks around 1.6s-2s (c) The time profile of Hb in the whisker barrel cortex. Soon after the stimulus, a slight peak in Hb can be observed at 0.6s. Hb follows a continous dip following that initial peak

### Percentage change in hemodynamic response at various points in the barrel sheet

The hemodynamic response to a whisker stimulus varies throughout the capillary sheet depending on which whisker is stimulated. We considered the total area of the sheet as 4*mm* X 4*mm*. The response of the entire sheet is observed after giving input stimulus at C3 whisker. The whisker barrel corresponding to the stimulated whisker is known as the principal barrel. Three points are identified in the capillary sheet which is aligned to the whisker barrel cortex, one at the centre of C3, another point at the boundary of C3 and D3 and one point at a long distance from centre of C3, at the centre of capillary area corresponding to E1 barrel. An area of dimension 300μm X300μm around the centre of principal barrel is considered as the region of interest (ROI). The average hemodynamic response (HbT) over approximately 300μ*m* X 300μ*m* area surrounding each of the three points is plotted in Fig.4.a. As seen by Devor et al(11), the peaking of HbT response in principal barrel occurs after around 2 seconds after the presentation of input stimulus. A decrease in HbT outside the region of interest is observed but did not match the experimental results suggesting additional vasoconstriction happening in the nearby vessels.

**Figure 4:**
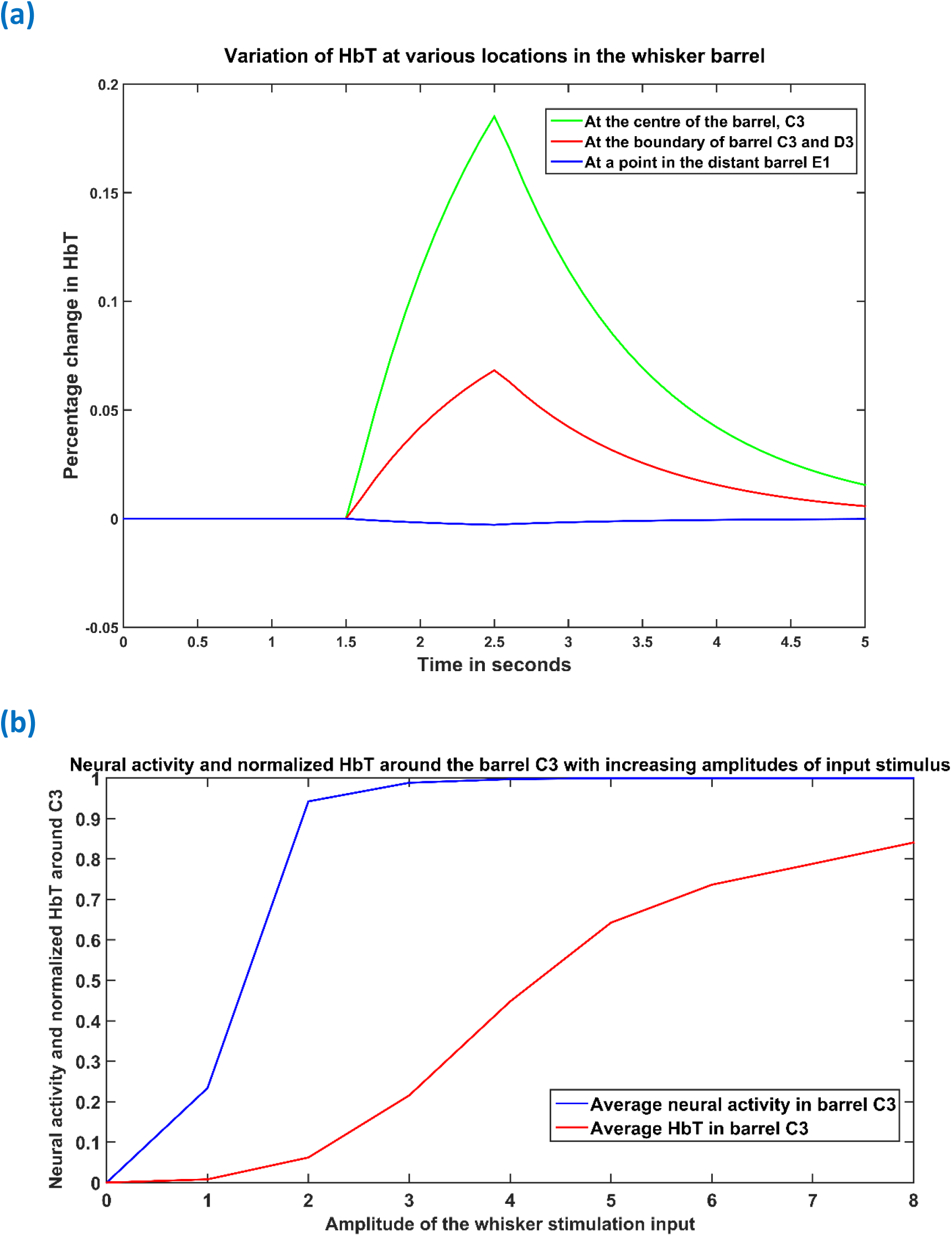
(a) Peak HbT observed 2 seconds post stimulus presentation to C3 whisker, (i) at the centre of the barrel corresponding to the whisker (principal barrel) which was stimulated (green), at the boundary of the principle barrel (red) and very far from the pricipal barrel (blue). (b) The average of peak response of HbT and the average neural activity around the principal barrel for different stimulus amplitudes

### Hemodynamic response increases beyond saturation of neural activity

The increase in neural response and hemodynamic response with increase in the amplitude of input stimulus does not follow the same pattern. Devor et al(11) observed that when the stimulus amplitude was increased continuously, the neural activity over an area saturates very soon, whereas the hemodynamic response kept increasing monotonically. We presented the network with input stimulations of various strengths defined by the amplitude of the Gaussian activity in the whisker pad. In Fig.4.b, the X-axis denotes the strength of input stimulus. The peak HbT is observed at an area around the principal barrel and the average of this peak HbT over that area is noted for each stimulus amplitude. The average neural activity is also noted around the same area for each stimulus amplitude. It is observed that the neural activity (blue) quickly reaches saturation whereas HbT (red) slowly increases with stimulus amplitude. The model is able to capture the saturation of the neural activity and the monotonous increase of hemodynamic response over an area while presented with stimuli of increasing amplitudes.

### Change in ATP at the tissues

The effect of vascular response to neural activity is reflected in the variation of ATP concentration in the neuron as shown in figure 5b. This pattern of change is similar to that observed by Ching et al(24). In order to blend an abstract model like LISSOM and a biophysical model like the vascular network, we consider the variable ATP in Eq.28 to be a dimensionless representation of ATP concentration. The threshold, which decides the firing ability of neuron (Eq.1) depends on this ATP concentration (Fig.5.a) according Eq.5. The idea of varying neural activation threshold as a function of ATP is consistent with the neurovascular model of Chhabria et al(8). The threshold and the ATP are dimensionless parameters to be included in the LISSOM equations. Fig. 5.b shows the change in ATP after the stimulus was given at t=1s. The initial dip followed by the slow rise is in agreement to the study by Chhabria et al(8). If the ATP variable decreases below the critical value (ATP=2 in this case), results in a high threshold of firing in the neurons reducing the firing probability.

**Figure 5:**
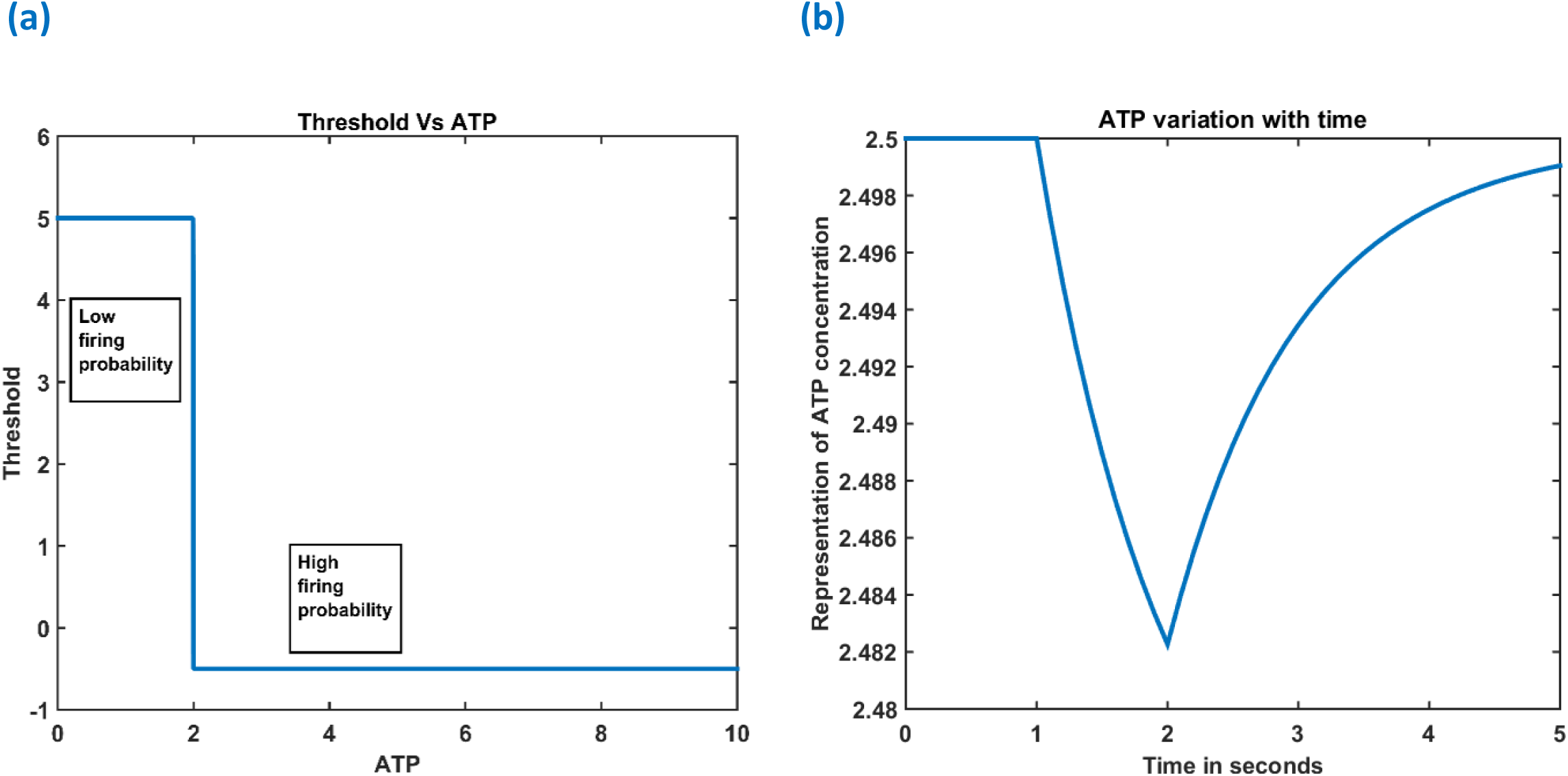
(A) Variation of threshold as a function of available ATP (B) Observed variation in ATP around the principal barrel post stimulus presentation (stimulus is given at t=1s)

### Influence of vascular pathology in whisker barrel map reorganization

In order to observe the influence of vascular pathology on whisker barrel map reorganization, the LISSOM network has to be retrained in the presence of the vascular feedback. Since the network with the whole whisker barrel is computationally intensive, a small area of whisker barrel cortex representing six whiskers, 3 whiskers each of C and D rows are considered (indicated by the red circle in Fig.9.b). The retraining in LISSOM is justified by considering the rats to be in early stages of development where plasticity is high. The neural layer gets retrained and the map reorganizes in case of any metabolic mismatch that causes a change in the neuronal response to the stimuli (20). The whisker barrel map reorganization is observed under four conditions: 1) Control, 2) Lesion, 3) Hypoxia –Lesion, and 4) Hypoxia-Ischemia as explained in the methods section. The plasticity of the network in whisker lesion conditions is observed by the increase in the ratio of the area in the whisker barrel cortex which represent D row to an area which represent C row (D/C ratio). Fig 6 (A-D) shows the whisker barrel reorganization under conditions 1 to 4. It can be observed that the area in the neural sheet representing C whiskers and D whiskers are almost equal when the retraining is done with a healthy vascular feedback (Fig 6.A). The lesioning of the C whisker lead to the reduction in area representing C whiskers and increase in the area representing D whiskers (Fig.6.B) displaying plasticity. Fig. 6.C shows that the plasticity is more or less preserved under hypoxic condition also as observed by Ranasinghe et al (20). The fourth condition which is the ischemia combined with hypoxia caused a global reduction in blood flow and due to inadequate feedback from the vessels, the neurons had a very low firing probability. This is due to the increase in the threshold value resulting from a low ATP value. Because of this low firing probability, plasticity does not take place and hence the response map remains unchanged as observed by Ranasinghe et al(20). The peak response to the stimulus is very low in the hypoxia ischemia condition due to large threshold value. Therefore, the response map shown in this case is not that of the output, but the net input obtained before passing through the sigmoid function. Basically, it is equivalent to displaying subthreshold activity of the neuron instead of the usual spiking response. This is similar to a situation where a neuron which is tuned to a particular whisker does receive the maximal input but due to the raised threshold, it is unable to fire. Because of this, the plasticity of the network is lost as observed in Fig. 6.D. The fraction of cells representing the D whiskers to C whiskers under all four conditions are compared with that of experimental observation in Fig. 7.a. The possible effect of stages of hypoxia ischemia and pure ischemia on plasticity as observed by the model is shown in Fig. 7.b. The effect of various stages of hypoxia as observed from the model is also plotted in Fig.7.c.

**Figure 6:**
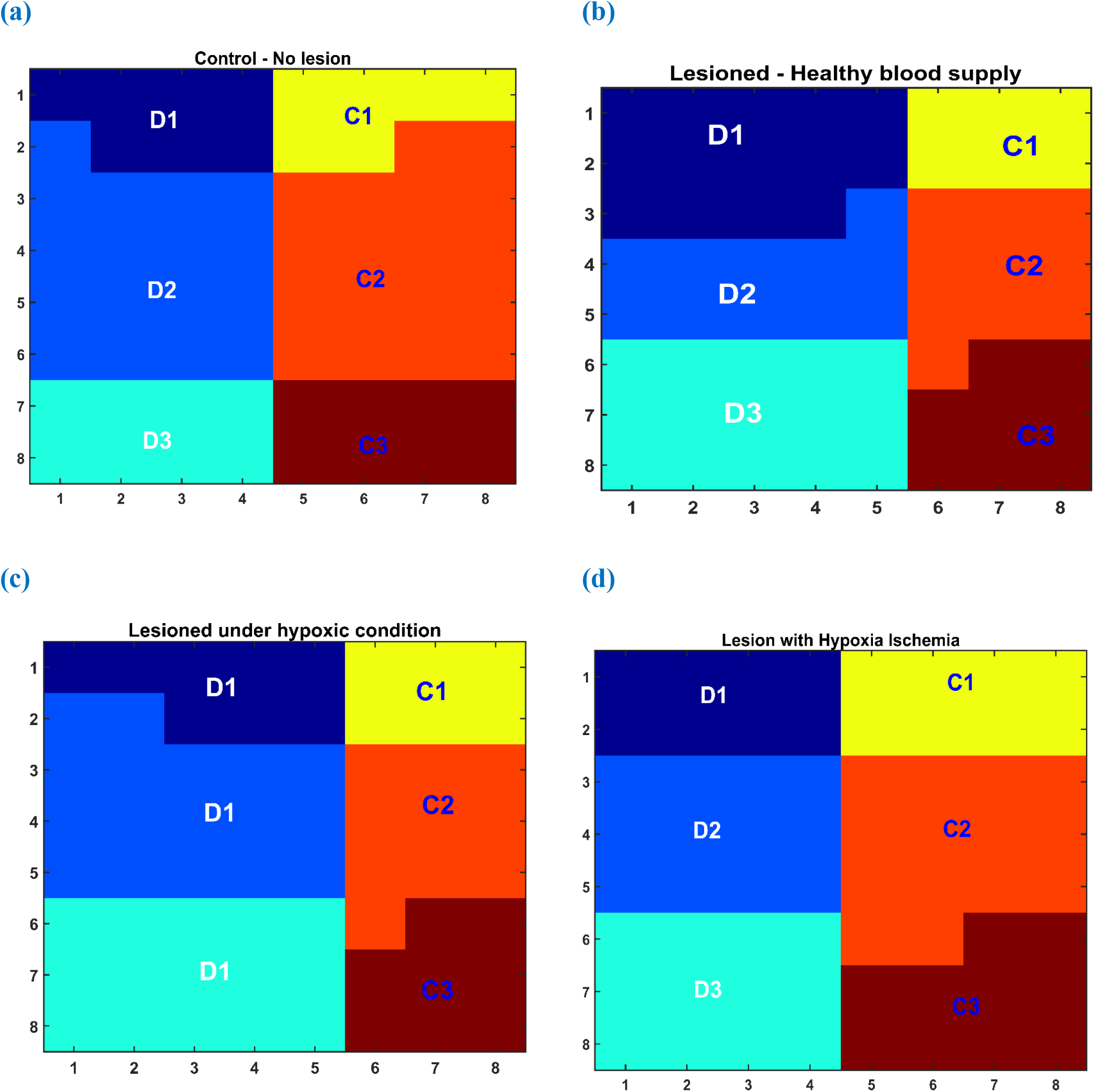
(A)-(D) Topographic map formation around a small area in whisker barrel cortex under various conditions. (A) When the whiskers are intact, and blood supply is normal (B) When the C1-C3 whiskers are lesioned, but blood supply is normal. (C) When the whiskers C1-C3 are lesioned and also the blood is hypoxic. (D) When the whiskers C1-C3 are lesioned and also under hypoxia ischemia condition

**Figure 7:**
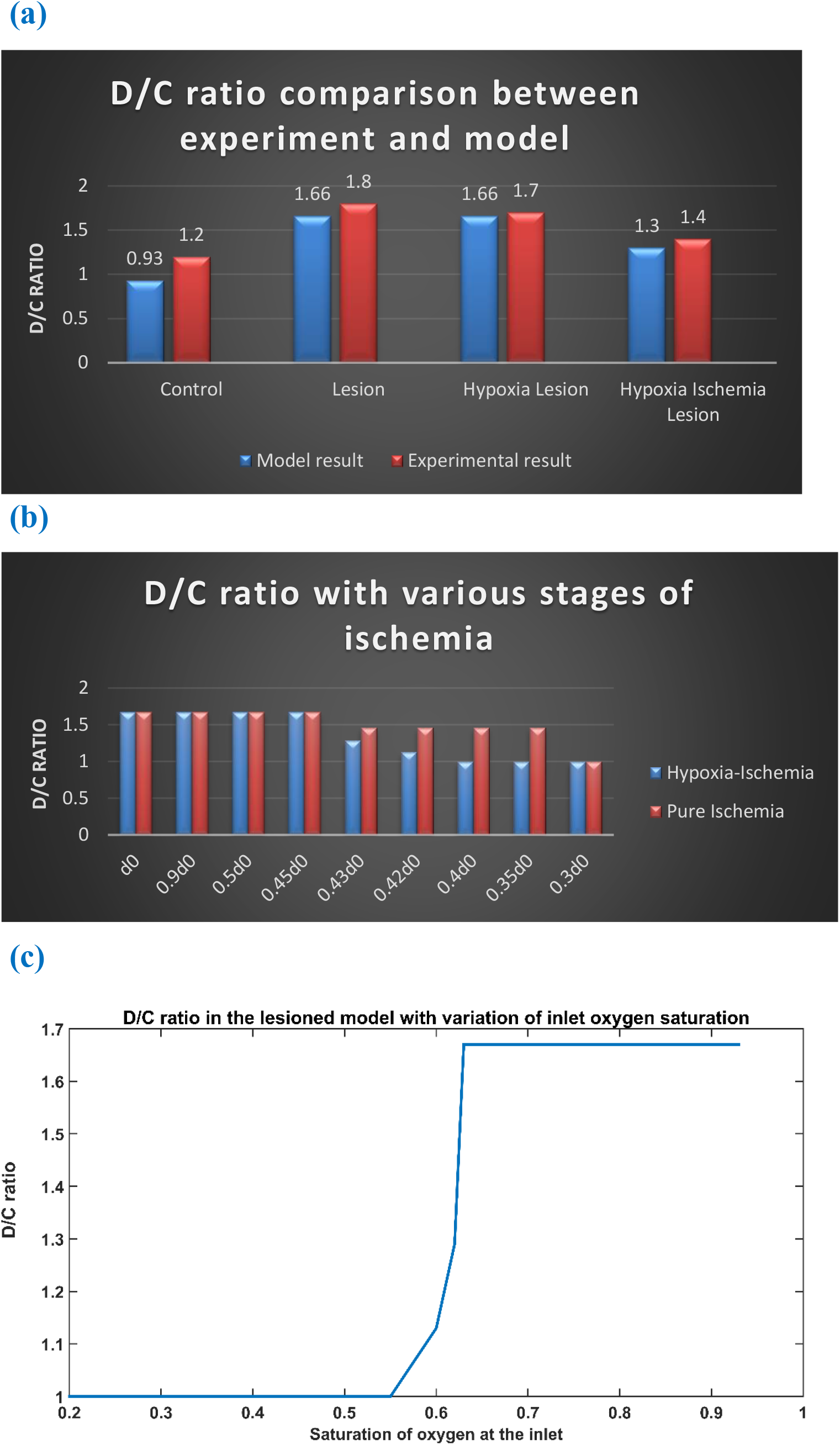
(a) Comparison of D/C ratio of experiment and model under the four simulation paradigms (b) The change in D/C ratio of a lesioned model at various stages of hypoxia-ischemia(blue) and purely ischemic condition (red) (c) The change in D/C ratio of a lesioned model at various percentages of oxygen saturation at the inlet.

## Discussion

A physical neural system can carry on with its firing activity only if it has an adequate and timely supply of energy. The vascular contribution to efficient neural performance has always been taken for granted. Our model is an attempt to capture the influence of retrograde signalling from vessels on the neural network characteristics. Our model captures the neurovascular coupling at a network level in rat whisker barrel cortex. It assumes a middle ground between detailed biophysical models(7,25,26) and abstract models(6)(9). The key feature of the model is that it captures the feedback from the vascular layer to the neural layer, representing the dependence of neural activity on the energy substrates released by the cerebral vessels. The behaviour of each neuron changes depending on the energy available to it and hence influences the network properties in a significant manner. Vascular structure, defined by a three-dimensional branching of blood vessels at different levels of hierarchy (arteries, arterioles, capillaries), perfuses a two-dimensional neural sheet at the level of capillaries to retrieve the experimentally observed hemodynamic responses(10,11,23) in the rat whisker barrel cortex. This model can be seen as a first step towards capturing the variations in hemodynamic responses observed in different areas of the brain(27). Individual units of LISSOM encode the rate of neuronal firing over an area. The forward connection to the vessels represents the vasodilatory influence of the neural firing and is represented by a variable that affects the elasticity of the vessel wall. The oxygen consumption near the tissues is directly dependent upon the neural activity and available oxygen, which is a direct indication of the metabolic demand. The feedback signal from the vessels to neurons is in the form of available oxygen for metabolism whose end result is ATP. The model treats the metabolic aspects of neurovascular interaction by considering first order dynamics for the cerebral metabolic rate of oxygen (CMRO_2_) with the synthesis term proportional to the net oxygen extracted from the vessels (OE) and the neural activity. ATP derived from this oxygen metabolism is again assumed to follow a first order dynamics with synthesis term being governed by CMRO_2_ and the decay being proportional to the neural activity. The equation predicts a dimensionless quantity (ATP) which replicates the characteristics of concentration of adenosine triphosphate([ATP]). The profiles of CMRO_2_ and [ATP] are in agreement with the experimental observations(28,29). The dimensionless parameter representing [ATP] could be incorporated easily into the LISSOM network made up of sigmoidal neurons by making the threshold of the neurons a function of the dimensionless variable, ATP.

This model which merges the biophysical metabolic quantities of vessels and abstract activation variables of the neural layer was able to reproduce the hemodynamic response observed experimentally. The time profiles of the hemodynamic responses (HbT, HbO and Hb) were in agreement with the experimental observations. By introducing the delay of vasodilatory signal in reaching the vessels, the model was able to capture the immediate drop in oxygenated haemoglobin observed in the proximal vessels of excited neurons soon after the stimulus is given. The model was also able to replicate the change in hemodynamic variables with spatial accuracy. It was able to replicate the vascular dilation/HbT variation at various locations on the whisker barrel after stimulating a single whisker. We can also control the amount of dilation in arterioles and capillaries individually to suit the literature (18).

As a first step to explore the influence of vascular feedback on neural network properties using this framework, we modelled the plasticity of the brain under conditions of hypoxia/ischemia in neonatal rats. The retraining of the LISSOM layer with vascular feedback was carried out for four different paradigms for four different vascular characteristics. Observing the response map of the retrained LISSOM network revealed the influence of vascular feedback on neural reorganization due to plasticity. Our model replicated the effects seen on neonatal rats until postnatal day 3(20). In their study, Ranasinghe et al showed that the neonatal rats had plasticity preserved as in controls, in hypoxic condition alone, but the plasticity was lost during hypoxia-ischemia condition. In the hypoxia-ischemia condition, the cortex was unable to respond to stimuli preventing any plasticity. The model supported the experimental observation that the hypoxic condition (oxygen saturation as low as 70%) alone does not affect the plasticity significantly. The model could also capture the absence of plasticity during Hypoxia-Ischemia condition. Thus, this experiment shows that short-term hypoxia does not compromise the neural functionality as compared to ischemia. But as observed in Fig. 7.c, a drop of inlet saturation below 65% does affect the plasticity. Similarly, there is a difference in how the plasticity is affected with reduction in arteriolar diameter under normal oxygen supply and hypoxic condition. To the best of our knowledge, a study on how a step by step reduction of blood flow or a step by step reduction in oxygen concentration has not yet been carried out. As seen in Fig.7.b and Fig.7.c, the model predicts that there is an intermediate stage where plasticity is not completely lost. This emphasises the need for an extensive experimental study where the neural responses are observed under carefully calibrated vascular changes.

As discussed by Hillmann(30), although a number of candidate mechanisms for neurovascular coupling have been identified, their integrated role has still not been understood. The vascular origin of a number of neurodegenerative diseases(31–34) points to the importance of understanding the proper functioning of neurovascular communication. Apart from this, the role of vessels in neural information processing is also a topic of debate(1,35). Vascular characteristics like vasomotion is an indication that vessels have their own dynamics which may influence each other and also the neuronal dynamics(35–37). With an intact network layer structure for neurovascular interaction, by simply modifying the vessel model to incorporate active and intrinsic dynamics, - and not just the vascular rhythms passively driven by neural inputs, - the effect of vascular dynamics on neural dynamics could be studied which is going to be the objective of the next version of our model. The individual unit of LISSOM network in this model is a sigmoid neuron. This could be replaced by a network of spiking neurons which represent neural activity more accurately. The vascular network in this model is a simple tree structure of variable resistors in which while the flow is simply analogous to the current flow, the saturation of oxygen in the blood is also given equal importance. Also, the neurovascular interaction could be mediated by modifiable weight connections which makes possible multiple vascular adaptations like arteriogenesis(38,39) and angiogenesis(40,41) following ischemic conditions..

One drawback of this model is that we did not consider the lateral interaction among vessels, which, we believe, would be crucial for optimal distribution of blood flow. We also have not explicitly considered the role of astrocytes in the neurovascular coupling which might play an important part in both feedforward and feedback interactions between neuron and vessels, even though the generalized vasodilatory parameter takes care of that factor to an extent. The generalized vasodilatory parameter acts on the capillaries which are mediated by astrocytes and on the arterioles, which are mediated directly by neurons (18).

## Material and Methods

The proposed model for bidirectional neurovascular interaction consists of a three-dimensional vascular network which branches from a large pial artery into smaller arterioles and capillaries to perfuse the two-dimensional neural sheet at the level of capillaries (fig 8.a). The whisker stimulations act as input stimuli to the neural layer and its response is conveyed to the vascular layer in the form of vasoactive signals causing vessel dilation/contraction. The vascular response determines the release of ‘energy’ to the neural layer, which controls the threshold of neural activation.

**Figure 8:**
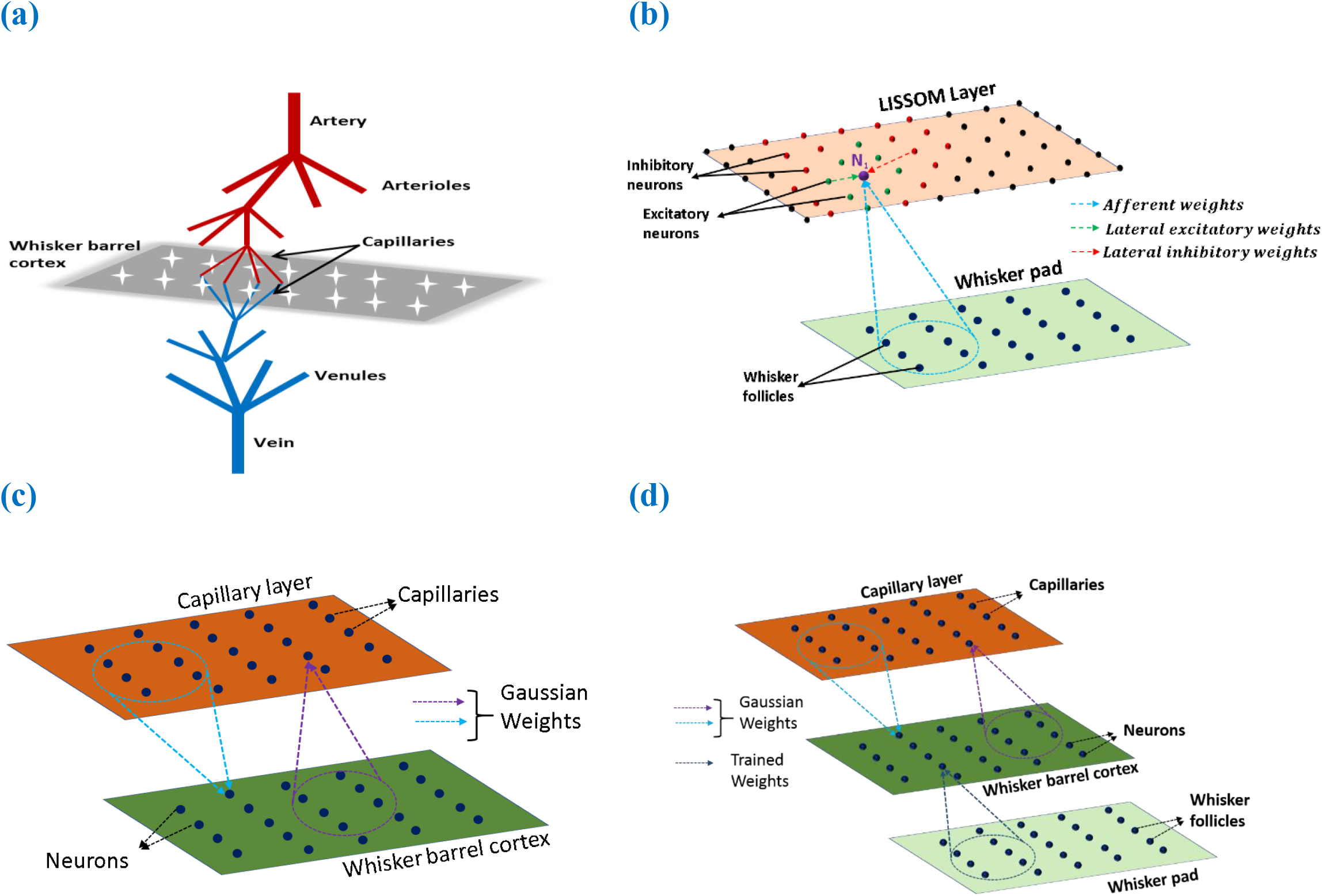
(a) A schematic of the model. The big pial artery branches to arterioles and they again branch to capillaries to perfuse the neurons. The neurons and vessels interact at the level of these capillaries. (b) LISSOM architecture. The neuron *N*_1_ receives weighted input from an area in the whisker pad (blue dots). It receives excitatory input from the immediate neighbouring neurons (green dots) and inhibitory input from distant neurons (red dots)(c) The neurovascular connectivity. Each vessel receives vasodilatory information from an area of neurons (violet dotted circle). Each neuron receives oxygen from a group of proximal vessels (blue dotted circle). (d) A schematic for complete interactions among the three layers, Input layer (whisker pad), Neural layer (whisker barrel cortex), and blood vessels (capillaries). The only trained weights are from input layer to whisker barrel cortex (dark blue dotted circle)

### Neural Layer

The neural layer, which represents the V4 layer of rat whisker barrel cortex (42), is modeled using a LISSOM, a biologically plausible model of cortical activity. The LISSOM is a two-dimensional network of neurons arranged on a grid as shown in (Fig.8.b). Each neuron (N_1_) is connected to a small region of the input known as receptive field using trainable afferent weights. Each neuron receives an afferent signal which is a weighted sum of the intensity values in the receptive field. The neurons are also laterally interconnected using trainable lateral weights in such a way that the neighbouring neurons within a specified radius (excitatory radius) of any given neuron (N_1_) gives excitatory input and those neurons which lie outside the excitatory radius but within a specified outer radius (inhibitory radius) of the neuron N_1_ gives inhibitory input to it. All the input signals are summed up and passed through a sigmoidal activation function. This center-surround pattern of lateral connections induces competition and helps in the formation of topographic maps as seen in the rat whisker barrel cortex.

The neuronal substrate for the LISSOM architecture is constrained so that the entire barrel cortex has a parabolic outline mimicking a real barrel cortex. The parabolic constraint is essential to produce barrels with a similar organization as in the real barrel cortex. Interestingly, a similar curvilinear boundary constraint was found to be critical to model retinotopic map formation(43). The constrained LISSOM, when trained on the whisker stimulation input, forms topographic maps similar to the whisker barrel map seen experimentally(42).

The whisker pad on the snout of rats is simulated by considering 24 whiskers arranged as shown in Fig (9.a)(21). Each blue dot represents one whisker and is identified using the row and column coordinates with numerals 0-4 representing column and A-E representing the rows. The stimulation on each whisker is modelled as a two-dimensional Gaussian function with peak at the location of the whisker being stimulated. The amplitude of the peak of Gaussian function defines the amplitude of the whisker stimulation.

**Figure 9:**
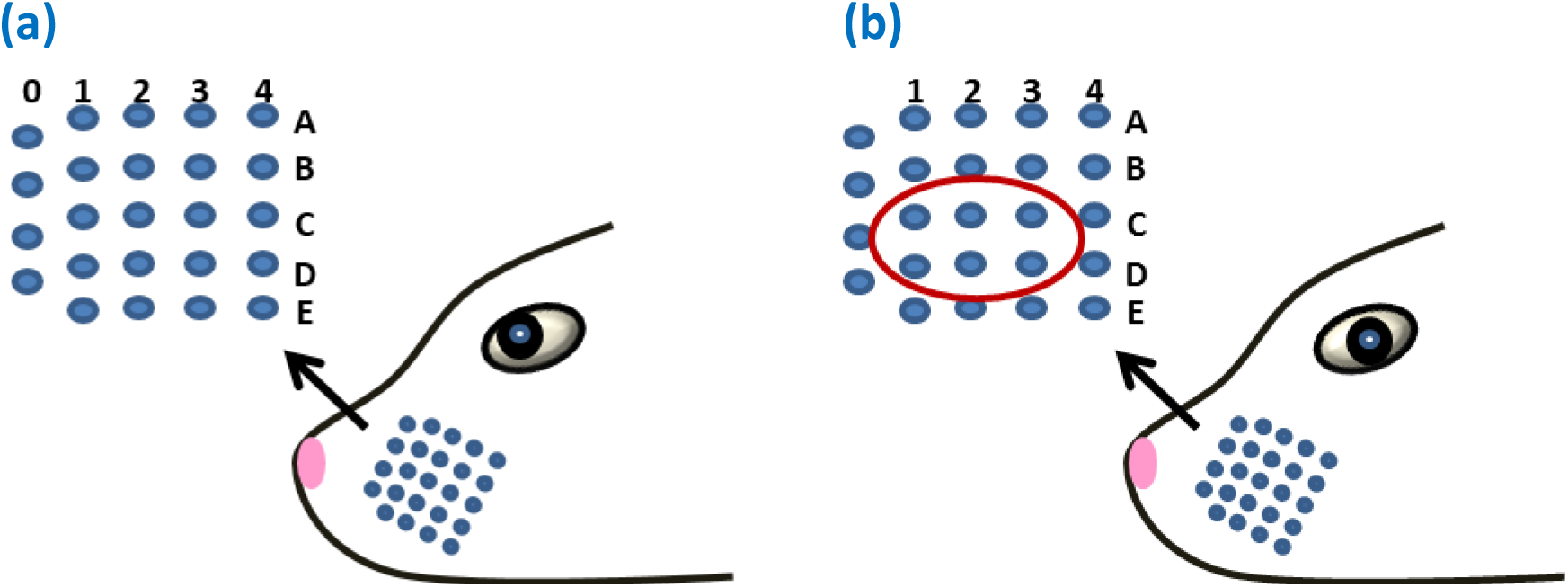
(a) The whisker pad of rat. Each whisker is addressed using the row (A-E) and the column (0-4) (b) The whiskers which are being considered for pathological study is shown in red circle (C1-C3 and D1-D3)

The vascular layer is modeled as a 3-dimensional tree structure with arteries branching first into penetrating arterioles and the arterioles branching into capillaries. The big pial artery gives rise to 16 smaller arteries and each of the arteries further branches to 16 arterioles giving rise to a layer of 256 penetrating arterioles. Each of the penetrating arterioles gives rise to 16 capillaries altogether forming a capillary bed with 4096 capillaries. The capillary bed aligns itself to the neural layer. Like in the case of neurons, each capillary is indexed by its location on the two dimensional grid (Fig.8.c). The assumption is that for every neuron, there exists a capillary. It is at the level of capillaries that the oxygen exchange takes place. In this model, we assume that the capillaries as well as arterioles dilate in response to the neural activity. Each capillary is connected to a small receptive field (perfusion field) in the neural layer bidirectionally by an untrained weight matrix defined by two dimensional Gaussian functions as shown in Fig.8.c. The Gaussian weight connection ensures that the neurons influence the nearest capillaries and in turn get maximum nourishment from the nearest vessels. One assumption that we make here is that each vessel (capillary) is influenced by a neighborhood of neurons and the radius of this neighborhood (receptive field/perfusion field) is taken as approximately half of the perfusion field observed for penetrating arterioles (400μmX400μm)(44) since we do not have exact biological values identified for the perfusion field of capillaries. The standard deviation of the gaussian weight connections are fixed in such a way that each capillary perfuses roughly 200μmX200μm.

Figure 10 shows the complete schematic of the interaction between neural and vascular layer. The neural activity modifies the elasticity factor (β) which changes the pressure-volume relationship in the vessel. This causes a redistribution of the blood flow resulting in dilation of some vessels and constriction of others. The change in flow rate and the volume (V) influences the amount of oxygen saturation (S) in the blood. The amount of oxygen that diffuses out of the capillaries (OE) would be dependent on the gradient of partial pressure of oxygen (*PO*_2_) in the vessels and the neural tissues. This oxygen which reaches the tissues influences the ATP production by oxidative phosphorylation indicated by cerebral metabolic rate of oxygen (CMRO_2_). The available ATP at the tissues in turn influences the threshold of neural firing.

**Figure 10:**
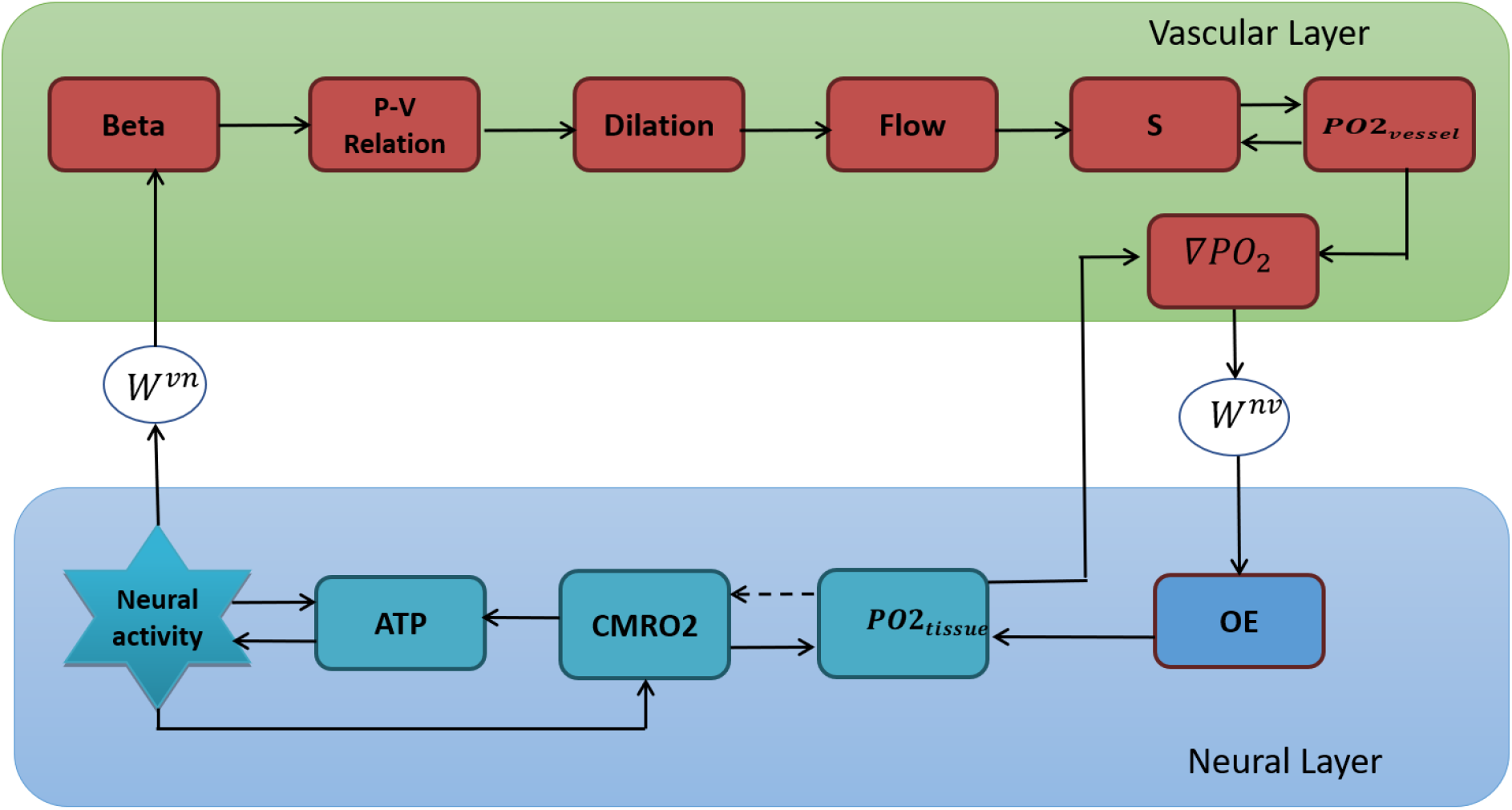
A schematic of the interactions among various quantities involved in neurovascular coupling. Beta (elasticity of the vessel, β). P (pressure), V (Volume), S (Saturation of oxygen in blood), PO2(partial pressure of oxygen), OE (oxygen extraction), CMRO2 (cerebral metabolic rate of oxygen), *W^vn^ and W^nv^* (Gaussian weights), ATP (adenosine tri phosphate).

### Modeling the neural response

The LISSOM sheet consists of *nxn* neurons each of which is indexed using the row and column information in the grid. The net input of each neuron is determined by a weighted contribution of afferent input, total excitatory input from the neurons in the excitatory radius and the inhibitory input from the neurons in the inhibitory radius. The output of each neuron is defined using Eq.1

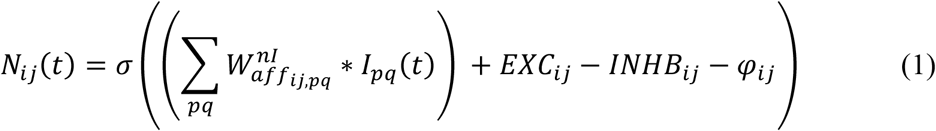

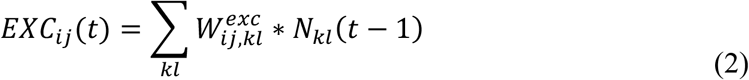

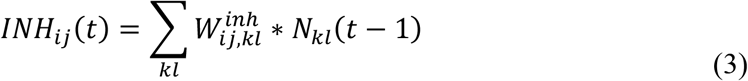

Where 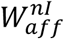 is the afferent weight stage from input layer I to neuronal layer n, *W^exc^* is the excitatory weight stage and *W^inh^* is the inhibitory weight stage. (*i, j*) denotes the index of the neuron N_1_ in the two-dimensional grid. (*k, l*) denotes the index of a neuron in the neighbourhood of N_1_ which gives excitatory or inhibitory projections to neuron N_1_ depending on their proximity to N_1_. *σ* is the activation function defined as,

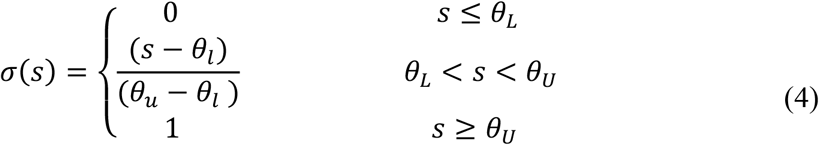

Where *θ_L_ and θ_U_* are the lower and upper thresholds of the sigmoid function. In eqn. (1), *φ_i,j_* is a threshold variable whose value depends on the available ATP near the neural tissues.

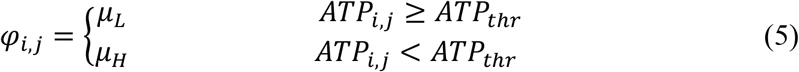

Where *ATP_thr_* is defined as the minimum ATP required for proper functioning of neurons. *μ_L_* and *μ_H_* are set in such a way that the *μ_H_*>*μ_L_*.

All weights are trained using Hebbian learning as shown in eqn. 6 for training lateral connections using asymmetric learning and eqn. 7 for afferent weight training

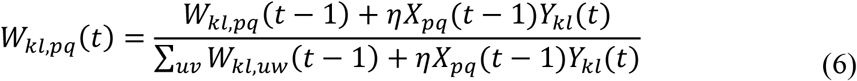

Where *W_kl,pq_* is the lateral connection (excitatory or inhibitory) from neuron (*p, q*) to neuron (*k, l*), *η* is the learning rate, *X_pq_*(*t* − 1) is the settled presynaptic neuron activity at instance *t* − 1 and *Y_kl_*(*t*) is the settled activity of neuron (*k, l*) at time *t*.

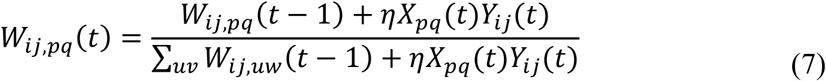

Where *W_ij,pq_* is the afferent connection weight from input pixel (*p, q*) to neuron (*i, j*), *η* is the learning rate, *X_pq_*(*t*) is the input at instance *t* and *Y_ij_*(*t*) is the settled activity of neuron (*i, j*) at time t.

### Modelling the vascular layer

Each blood vessel is considered to be a cylinder with diameter(d) and length(l) depends on the level of branching from the pial artery. The values of the diameter and length are adopted from Boas et al(17). The other important variables that define the vessel characteristics are the Resistance (R), Volume (V), Pressure at the centre of each vessel (*P_e_*), Pressure at the node (*P_n_*) from which branching of vessels take place, the elasticity of the vessel (*β*) and Saturation of oxygen in blood (S). Each capillary at the grid location (i,j) is charecterised by the length (*l_i,j_*), resistance (*R_i,j_*) and diameter (*d_i,j_*).

The resistance of each vessel can then be calculated using Poiseuille’s law,

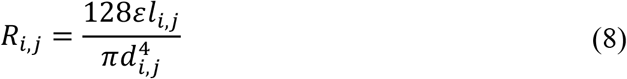

where *ε* is the fluid viscosity. The neurovascular coupling arises from the elasticity factor *β* which is a function of neural activity. This factor results in the vasodilation of the proximal vessels. The various pathways that contribute to the vasodilation are considered by incorporating a time delay for the neural influence on the elasticity factor.

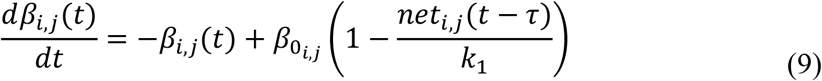

where *net* ∈ [0,1] is the weighted sum of neural activity (N) over an area in the LISSOM at time (*t* − *τ*), *β*_0_ is the *β* at resting state and *k*_1_ is a constant.

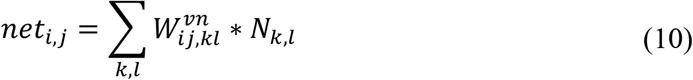

Where 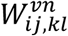 is the weight of the connection from the neuron at location (*k, l*) in the neural layer to the vessel at location (*i, j*) in the vascular layer. The strength of the weight from a neuron (*k, l*) to a vessel (*i, j*) drops with its distance from the vessel. Hence the weight matrix projecting to a vessel from an area in neural layer is defined by a Gaussian weight distribution centred at the coordinate of that vessel. The weight of connection from a neuron (*k, l*), to a vessel (*i, j*) is defined by

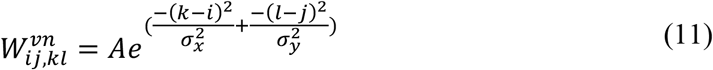

Where A is the amplitude of the Gaussian, and *σ_x_* and *σ_y_* are the standard deviations. We allowed the dilation of a given penetrating arteriole to depend on an average of the effective neural activity felt by the branching capillaries originating from that arteriole.

The pressure at the center of the vessel (*P_e_*) and volume (V) of the vessel are assumed to be related linearly as follows,

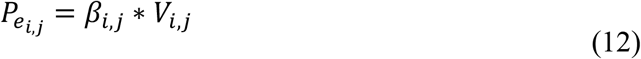

An increase in the neural activity causes a pressure drop in the proximal vessels enabled by a reduction in the elasticity factor *β*. The pressure drop caused at edge(s) due to the neural activity causes a redistribution of nodal pressures. The redistributed pressure is calculated by rewriting flow balance at a node in terms of nodal pressures and edge presures.

The change in pressure, in turn, redistributes the flow of blood into the vessel and out of the vessel, building up the volume in some vessels. The flow of blood in and out of the vessels is calculated as follows.

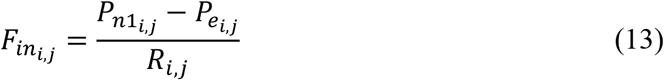

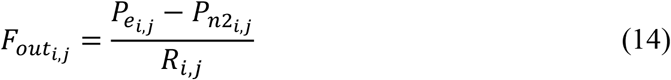

At rest, *F_in_* = *F_out_*. Neural activity induced changes in pressure disturbs this equality causing vasodilation or constriction. This causes a change in the volume given by.

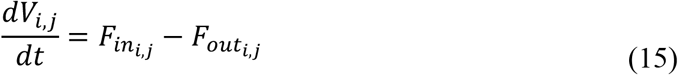

This change in volume and flow rate causes a change in oxygen saturation in the vessels. The rate of change of oxygen saturation depends on the difference in flow rate at the inlet and outlet at each vessel, the amount of oxygen extracted from the vessel (*OE^v^*), and the volume of the vessel. The saturation at the inlet of any vessel is assumed to be the saturation at the node from which it branches. The saturation at the outlet is again assumed to be the same as the saturation at the node to which it is connected at the bottom. The nodal saturations are calculated using the oxygen flow balance equation. At any node, the mass is conserved: the products of flow and saturation terms should add up to zero. The saturation of blood depends on the direction of current flow. It is crucial to identify the immediate source point of flow to get a balanced mass-flow condition at each node.

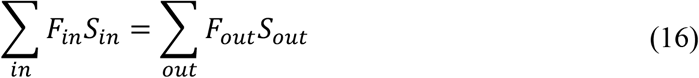

The rate of change of saturation is given by the following relationship derived from Boas et al(17).

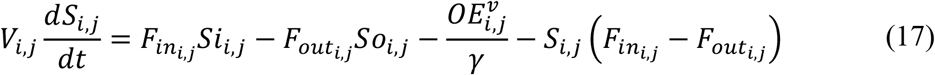

Where *γ* is a constant representing the concentration of hemoglobin. Once the saturation of hemoglobin (S) and the volume (V) are estimated, the concentration of total hemoglobin (HbT), oxygenated hemoglobin (HbO) and deoxygenated hemoglobin (Hb) an be calculated as follows.

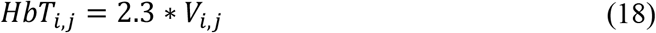

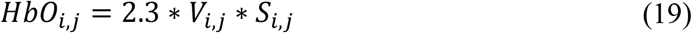

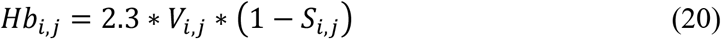

The change in oxygen saturation in a vessel brings about a change in the partial pressure of oxygen (*PO*_2_) in that vessel. The empirical relation(17) between the oxygen saturation and partial pressure of oxygen is assumed to follow

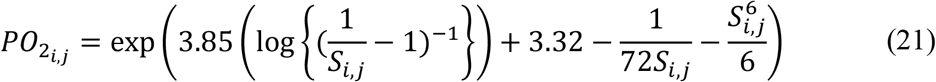

At the same time, the neural activity consumes oxygen in the tissues in order to avail energy via oxydative phosphorylation. The change in the metabolic rate of oxygen (*CMRO*_2_) at the tissue is normally estimated as a function of flow rate, oxygen extraction fraction and the hemoglobin concentration(45). But from the discussions in(46) and (47), neural activity appears to be a more direct correlate of CMRO_2_. Considering the fact that the total oxygen flux also has an equal significance, CMRO_2_ is influenced by neural signal and vascular signal equally. Thus it appears to be the key variable in modulating the feedback from vessels. In order to independently incorporate the effect of feedback from vessel to the tissues, based on the available oxygen at its neighbourhood, we suggest the following equation for the metabolic rate.

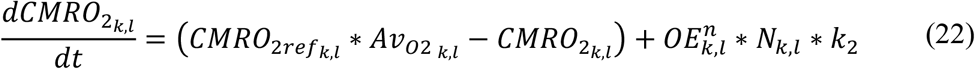

where *k*_2_ is a constant and *Av*_*O*2_ represents a fraction of available oxygen in the tissues, which is calculated as a fraction of *PO*_2*tiss*_ (partial pressure of oxygen) available at the tissues to the resting value of *PO*_2_ at the tissue (*PO*_2*tiss_max*_).

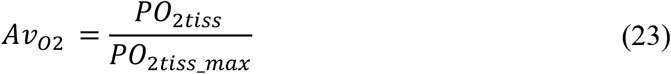

The equation (22) describes CMRO_2_ changes in a manner similar to the results observed in the studies by Boas et al(48), Jone et al(49) andMathias et al(26).

The increased oxygen metabolism results in a dip in the partial pressure of oxyen at the tissues. The rate of change of partial pressure of oxygen at the tissues depends on the total oxygen flux to the tissues (*OE^n^*) from the vessels and the rate of oxygen metabolism (CMRO_2_)

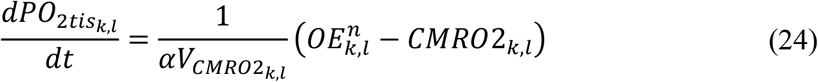

Where *αV*_*CMRO*2_ is a constant which represents the volume of tissue where the extracted oxygen is consumed. A reduction in the partial pressure of oxygen at the tissue leads to an increase in the extraction of oxygen from the vessels. The partial pressure of oxygen at the tissues as felt by each vessel is taken as a weighted sum of the partial pressure of oxygen at a small neural area around each capillary, corresponding to the receptive field of the capillary. The weight matrix used is the same Gaussian weight matrix used for calculating the effective neural activity at each vessel.

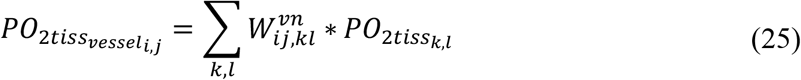

where *PO*_2*tiss_vessel_ij_*_ denotes the partial pressure of oxygen at the tissues as seen by the vessel at (*i, j*). The gradient between the partial pressure of oxygen at the capilliaries and the tissues lead to diffusion of oxygen from the capilliaries to the tissues, and is defined by the following equation.

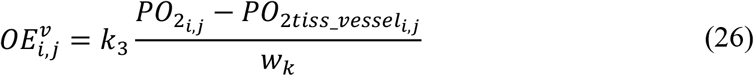

where k_3_ is a constant. The oxygen extracted from the vessels (*OE_v_*) diffuses to the tissues. The oxygen flux into each neuron is calculated as the weighted sum of the oxygen extracted from an area of proximal capilliaries which cater to the neuron. We assume here that most of the oxygen required by the neurons is exchanged at the level of capilliaries. The oxygen extracted by each neuron is given by,

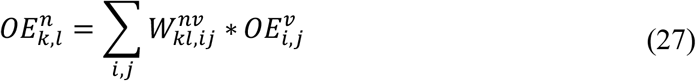

where 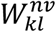 a gaussian weight distribution centered on the neuron at (*k, l*) similar to 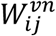 defined in eqn (11).

ATP is required to fuel neural firing activity. Most of the ATP available at the neural tissues is a result of oxidative metabolism(50). The cerebral metabolic rate (CMRO_2_), therefore, could be taken as an indication of ATP generation and the neural activity could be taken as a measure of ATP consumption. The relation between generation and consumption of ATP is assumed to be as shown below. ATP_ref_ is the resting state level of ATP at the tissues, and the CMRO2_ref_ is the cerebral metabolic rate during the resting state.

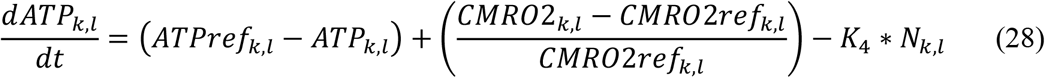

Where K4 is a constant. The first term ensures that the ATP values return to their resting values when there is no production and consumption. The second term indicates the net increase in ATP by considering the production of ATP proportional to the percentage of increase in CMRO_2_. The third term indicates the consumption of ATP proportional to the neural activity.

The above equations are solved using DDE23 and Euler method in MATLAB to find the hemodynamic responses to neural activity.

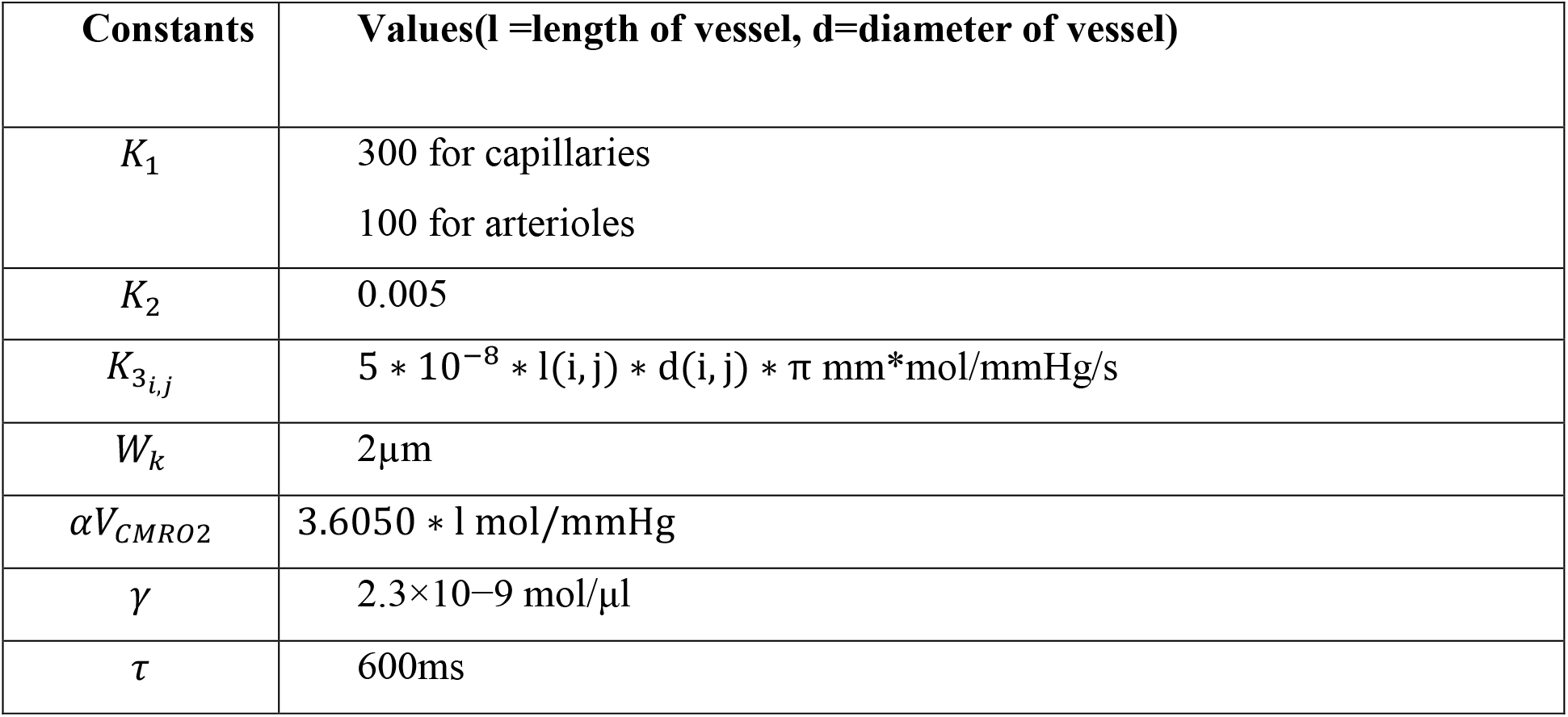

### The training paradigm

The LISSOM network is trained using the whisker stimulation input (Gaussian activation centred at each whisker location) for 4 epochs. The trained LISSOM is then connected to the vascular network forming three layered structure with input as the first layer, LISSOM as the second layer and Vascular bed as the third layer as shown in Fig.8.d. In order to check the hemodynamic response, the stimulus is given at time t=0 and retained for 1 second. The hemodynamic response is observed for 4s.

Simulating the network with all 24 whiskers is computationally very expensive. Hence for modelling pathological conditions, we consider a smaller network where input layer is a portion of whisker pad with just 6 whiskers. The neural layer with dimension 8×8 represents three barrels each of C and D row whiskers (fig.9.b). This is fed by a capillary bed of size 8×8. Once the pathology is introduced, the afferent connections from input layer to neural layer are retrained to observe the influence of the pathological condition in topographic map formation. The retraining of the LISSOM network is carried out in the presence of vascular feedback under four conditions: i) Control ii) Lesion iii) Hypoxia –Lesion iv) Hypoxia-Ischemia. In the control condition, the network is retrained in the presence of healthy vascular feedback using the input stimuli from all the six whiskers. In the lesioned condition, one row of the whisker is assumed to be lesioned and hence retraining is carried out with just the stimuli from intact whiskers. The vascular feedback is healthy in this condition. In the model, we assume that the C whiskers are lesioned (the whiskers C1,C2 and C3 in fig.9.b). Hence, we train the network only by using the stimulations of D whiskers. The hypoxic condition is simulated by limiting the inlet oxygen saturation to 70% as compared to the 94% in the healthy controls(51). The study is also carried out under a range of oxygen saturation from 30% inlet oxygen saturation to 94% inlet oxygen saturation. The retraining of the LISSOM layer is carried out using input stimulations of D whiskers, under the assumption that C whiskers are lesioned. The final condition is that the arterial ligation is carried out to simulate ischemia along with hypoxia. We simulate it by reducing the diameter of the Pial artery along with reducing the inlet oxygen saturation to 70%. The retraining of LISSOM network is carried out using only D whisker stimulation input. The stimulation is given for 2 seconds preceded and succeeded by 1 second of rest period before giving the next stimulus. The training is run for 4 epochs for all 4 conditions. In order to plot the response map for all the four conditions, the response of each neuron is observed before passing through the sigmoid function. The neuron is assigned the label of that input to which it shows the maximal response.

## Code accessibility

All codes are available in the ModelDB repository (http://senselab.med.yale.edu/ModelDB/showModel.cshtml?model=247661) and access code will be provided on request.

## Acknowledgements

We thank The Department of Biotechnology (DBT), Ministry of Science and Technology, Government of India for funding this project (BIO/17-18/303/DBTX/SRIN). We also thank Dr. Ryan T. Philips and Ms. Karishma Chhabria for the valuable discussions.

## Contributions

BSK did model designing, coding, analysis of the results, and manuscript preparation. AK did model designing and coding. VSC did model designing, analysis of the results, and manuscript preparation. SP did model designing and manuscript preparation. All authors discussed the results and commented on the manuscript.

## Corresponding authors

Correspondence to Srinivasa Chakravarthy

## References

1. Moore CI, Cao R. The hemo-neural hypothesis: on the role of blood flow in information processing. J Neurophysiol. 2008;99:2035–47.

2. Sirotin YB, Das A. Anticipatory haemodynamic signals in sensory cortex not predicted by local neuronal activity. Nature. 2009;457(7228):475–9.

3. Leithner C, Royl G. The oxygen paradox of neurovascular coupling. J Cereb Blood Flow Metab. 2014 Jan 23;34(1):19–29.

4. Filosa JA, Bonev AD, Straub S V., Meredith AL, Wilkerson MK, Aldrich RW, et al. Local potassium signaling couples neuronal activity to vasodilation in the brain. Nat Neurosci. 2006;9(11):1397–403.

5. Kim KJ, Ramiro Diaz J, Iddings JA, Filosa JA. Vasculo-Neuronal Coupling: Retrograde Vascular Communication to Brain Neurons. J Neurosci. 2016;36(50):12624–39.

6. Gandrakota R, Chakravarthy VS, Pradhan RK. A model of indispensability of a large glial layer in cerebrovascular circulation. Neural Comput. 2010;22(4):949–68.

7. Chander BS, Chakravarthy VS. A Computational Model of Neuro-Glio-Vascular Loop Interactions. PLoS One. 2012;7(11):1–11.

8. Chhabria K, Chakravarthy VS. Low-dimensional models of “neuro-glio-vascular unit” for describing neural dynamics under normal and energy-starved conditions. Front Neurol. 2016;7(MAR):1–20.

9. Philips RT, Chhabria K, Chakravarthy VS. Vascular Dynamics Aid a Coupled Neurovascular Network Learn Sparse Independent Features: A Computational Model. Front Neural Circuits. 2016;10(February).

10. Devor A, Dunn AK, Andermann ML, Ulbert I, Boas DA, Dale AM. Coupling of total hemoglobin concentration, oxygenation, and neural activity in rat somatosensory cortex. Neuron. 2003;39(2):353–9.

11. Devor A, Ulbert I, Dunn AK, Narayanan SN, Jones SR, Andermann ML, et al. Coupling of the cortical hemodynamic response to cortical and thalamic neuronal activity. Proc Natl Acad Sci. 2005;102(10):3822–7.

12. Devor A, Tian P, Nishimura N, Teng IC, Hillman EMC, Narayanan SN, et al. Suppressed Neuronal Activity and Concurrent Arteriolar Vasoconstriction May Explain Negative Blood Oxygenation Level-Dependent Signal. J Neurosci. 2007;27(16):4452–9.

13. Dunn AK, Devor A, Dale AM, Boas DA. Spatial extent of oxygen metabolism and hemodynamic changes during functional activation of the rat somatosensory cortex. Neuroimage. 2005;27(2):279–90.

14. Martin C, Martindale J, Berwick J, Mayhew J. Investigating neural-hemodynamic coupling and the hemodynamic response function in the awake rat. Neuroimage. 2006;32(1):33–48.

15. Gao YR, Ma Y, Zhang Q, Winder AT, Liang Z, Antinori L, et al. Time to wake up: Studying neurovascular coupling and brain-wide circuit function in the un-anesthetized animal. Neuroimage. 2017;153:382–98.

16. Blinder P, Tsai PS, Kaufhold JP, Knutsen PM, Suhl H, Kleinfeld D. The cortical angiome: An interconnected vascular network with noncolumnar patterns of blood flow. Nat Neurosci. 2013;16(7):889–97.

17. Boas DA, Jones SR, Devor A, Huppert TJ, Dale AM. A vascular anatomical network model of the spatio-temporal response to brain activation. Neuroimage. 2008;40(3):1116–29.

18. Biesecker KR, Srienc AI, Shimoda AM, Agarwal A, Bergles DE, Kofuji P, et al. Glial Cell Calcium Signaling Mediates Capillary Regulation of Blood Flow in the Retina. J Neurosci. 2016 Sep 7;36(36):9435–45.

19. Miikkulainen R, Bednar JA, Choe Y, Sirosh J. Computational maps in the visual cortex. Computational Maps in the Visual Cortex. 2005. 1–538 p.

20. Ranasinghe S, Or G, Wang EY, Ievins A, McLean MA, Niell CM, et al. Reduced Cortical Activity Impairs Development and Plasticity after Neonatal Hypoxia Ischemia. J Neurosci. 2015;35(34):11946–59.

21. Chen-Bee CH, Zhou Y, Jacobs NS, Lim B, Frostig RD. Whisker array functional representation in rat barrel cortex: transcendence of one-to-one topography and its underlying mechanism. Front Neural Circuits. 2012;6.

22. Buxton RB, Uludağ K, Dubowitz DJ, Liu TT. Modeling the hemodynamic response to brain activation. Neuroimage. 2004;23(SUPPL. 1):220–33.

23. Berwick J, Johnston D, Jones M, Martindale J, Martin C, Kennerley AJ, et al. Fine Detail of Neurovascular Coupling Revealed by Spatiotemporal Analysis of the Hemodynamic Response to Single Whisker Stimulation in Rat Barrel Cortex. J Neurophysiol. 2008;99(2):787–98.

24. Ching S, Purdon PL, Vijayan S, Kopell NJ, Brown EN. A neurophysiological-metabolic model for burst suppression. Proc Natl Acad Sci. 2012;109(8):3095–100.

25. Mathias EJ, Plank MJ, David T. A model of neurovascular coupling and the BOLD response: PART I. Comput Methods Biomech Biomed Engin. 2017;20(5):508–18.

26. Mathias EJ, Plank MJ, David T. A model of neurovascular coupling and the BOLD response PART II. Comput Methods Biomech Biomed Engin. 2017;20(5):519–29.

27. Lecrux C, Hamel E. Neuronal networks and mediators of cortical neurovascular coupling responses in normal and altered brain states. Philos Trans R Soc B Biol Sci. 2016;371(1705).

28. Hoge RD, Atkinson J, Gill B, Crelier GR, Marrett S, Pike GB. Linear coupling between cerebral blood flow and oxygen consumption in activated human cortex. Proc Natl Acad Sci. 1999 Aug 3;96(16):9403–8.

29. Aubert A, Costalat R. A model of the coupling between brain electrical activity, metabolism, and hemodynamics: Application to the interpretation of functional neuroimaging. Neuroimage. 2002;17(3):1162–81.

30. Hillman EMC. Coupling Mechanism and Significance of the BOLD Signal: A Status Report. Annu Rev Neurosci. 2014;37(1):161–81.

31. Iadecola C, Davisson RL. Hypertension and Cerebrovascular Dysfunction. Cell Metab. 2008;7(6):476–84.

32. Kisler K, Nelson AR, Rege S V., Ramanathan A, Wang Y, Ahuja A, et al. Pericyte degeneration leads to neurovascular uncoupling and limits oxygen supply to brain. Nat Neurosci. 2017 Jan 30;20(3):406–16.

33. Iadecola C. The Neurovascular Unit Coming of Age: A Journey through Neurovascular Coupling in Health and Disease. Neuron. 2017 Sep;96(1):17–42.

34. Tarantini S, Tran CHT, Gordon GR, Ungvari Z, Csiszar A. Impaired neurovascular coupling in aging and Alzheimer’s disease: Contribution of astrocyte dysfunction and endothelial impairment to cognitive decline. Exp Gerontol. 2017;94:52–8.

35. Pradhan RK, Chakravarthy VS. Informational dynamics of vasomotion in microvascular networks: A review. Acta Physiol. 2011;201(2):193–218.

36. Stergiopulos N, Porret C a, De Brouwer S, Meister JJ. Arterial vasomotion: effect of flow and evidence of nonlinear dynamics. Am J Physiol. 1998;274(6 Pt 2):H1858–64.

37. Secomb TW, Pries AR. Information Transfer in Microvascular Networks. Microcirculation. 2002;9(5):377–87.

38. Scholz D, Cai WJ, Schaper W. Arteriogenesis, a new concept of vascular adaptation in occlusive disease. Angiogenesis. 2001;4(4):247–57.

39. Buschmann I, Schaper W. The pathophysiology of the collateral circulation (arteriogenesis). J Pathol. 2000 Feb 1;190(3):338–42.

40. Krupinski J, Kaluza J, Kumar P, Kumar S, Wang JM. Role of angiogenesis in patients with cerebral ischemic stroke. Stroke. 1994 Sep 1;25(9):1794–8.

41. Beck H, Plate KH. Angiogenesis after cerebral ischemia. Acta Neuropathol. 2009 May 14;117(5):481–96.

42. Petersen CCH. The functional organization of the barrel cortex. Neuron. 2007;56(2):339–55.

43. Philips RT, Chakravarthy VS. A Global Orientation Map in the Primary Visual Cortex (V1): Could a Self Organizing Model Reveal Its Hidden Bias? Front Neural Circuits. 2017;10(January).

44. O’Herron P, Chhatbar PY, Levy M, Shen Z, Schramm AE, Lu Z, et al. Neural correlates of single-vessel haemodynamic responses in vivo. Nature. 2016;534(7607):378–82.

45. Buxton RB, Frank LR. A model for the coupling between cerebral blood flow and oxygen metabolism during neural stimulation. J Cereb Blood Flow Metab. 1997;17(1):64–72.

46. Jeffrey K. Thompson, Matthew R. Peterson RDF. Single-Neuron Activity and Tissue Oxygenation in the Cerebral Cortex. Sci (New York, NY). 2003;

47. Chong SP, Merkle CW, Leahy C, Srinivasan VJ. Cerebral metabolic rate of oxygen (CMRO_2) assessed by combined Doppler and spectroscopic OCT. Biomed Opt Express. 2015;6(10):3941.

48. Boas DA, Strangman G, Culver JP, Hoge RD, Jasdzewski G, Poldrack RA, et al. Can the cerebral metabolic rate of oxygen be estimated with near-infrared spectroscopy? Phys Med Biol. 2003;48(15):2405–18.

49. Jones M, Berwick J, Johnston D, Mayhew J. Concurrent optical imaging spectroscopy and laser-Doppler Flowmetry: The relationship between blood flow, oxygenation, and volume in rodent barrel cortex. Neuroimage. 2001;13(6):1002–15.

50. Lin A-L, Fox PT, Hardies J, Duong TQ, Gao J-H. Nonlinear coupling between cerebral blood flow, oxygen consumption, and ATP production in human visual cortex. Proc Natl Acad Sci. 2010;107(18):8446–51.

51. Sicard KM, Duong TQ. Effects of hypoxia, hyperoxia, and hypercapnia on baseline and stimulus-evoked BOLD, CBF, and CMRO2in spontaneously breathing animals. Neuroimage. 2005;25(3):850–8.

